# On the edge: Evolution of polarity protein BASL and the capacity for stomatal lineage asymmetric divisions

**DOI:** 10.1101/2021.08.05.455184

**Authors:** Ido Nir, Gabriel O. Amador, Yan Gong, Nicole K. Smoot, Le Cai, Hagai Shohat, Dominique C. Bergmann

## Abstract

Asymmetric and oriented stem cell divisions enable the continued production of patterned tissues. The molecules that guide these divisions include several “polarity proteins” that are localized to discrete plasma membrane domains, are differentially inherited during asymmetric divisions, and whose scaffolding activities can guide division plane orientation and subsequent cell fates. In the stomatal lineages on the surfaces of plant leaves, asymmetric and oriented divisions create distinct cell types in physiologically optimized patterns. The polarity protein BASL is a major regulator of stomatal lineage division and cell fate asymmetries in Arabidopsis, but its role in the stomatal lineages of other plants is unclear. Here, using phylogenetic and functional assays, we demonstrate that BASL is a eudicot-specific polarity protein. Among dicots, divergence in BASL’s roles may reflect some intrinsic protein differences, but more likely reflects previously unappreciated differences in how asymmetric cell divisions are employed for pattern formation in different species. This multi-species analysis therefore provides insight into the evolution of a unique polarity regulator and into the developmental choices available to cells as they build and pattern tissues.

**HIGHLIGHTS:**

- BASL is a eudicot-specific regulator of stomatal lineage asymmetric cell divisions
- BASL protein evolution includes stepwise addition of polarity domains to an ancestral MAPK-binding chassis
- Cellular quiescence and BASL-guided polarity generate proper stomatal spacing in tomato
- Cell size and fate asymmetries are uncoupled in the tomato stomatal lineage

## INTRODUCTION

Patterned surfaces adorn representatives of all major clades of multicellular life. These patterns serve functional roles in mature organisms and can bear imprints of the developmental trajectories underlying their production. Asymmetric and oriented stem cell divisions are often employed in the creation and pattern of tissues, particularly when development is flexible and environmentally responsive. In plants, stomatal lineages on the surfaces of leaves are prime models for investigating how cell polarity and asymmetric cell divisions (ACDs) link to tissue-wide patterns (Lee and Bergmann, 2019).

In the stomatal lineage of the dicot angiosperm *Arabidopsis thaliana*, BREAKING OF ASYMMETRY IN THE STOMATAL LINEAGE (BASL) is the central regulator of cell size and fate asymmetries (Dong *et al*., 2009; Zhang et al., 2016b; Zhang *et al*., 2015). Asymmetric “entry” divisions among protodermal cells initiate the stomatal lineage (Figure 1A, stage II). During these entry divisions, BASL becomes polarly localized in a cortical crescent (Figure 1A, stage I-III) that guides the division plane to create cells of different size and identity, and ensures its own asymmetric inheritance (Figure 1A, stage III). Fate asymmetry between the smaller, meristemoid (M) daughter and larger, BASL-inheriting stomatal lineage ground cell (SLGC) daughter is critical for proper patterning of the leaf epidermis as the subsequent division types available to these cells differ (Figure 1A, stage IV). Through continued “amplifying divisions” in a spiral pattern, meristemoids can surround themselves with a buffer zone of non-meristemoid cells before differentiating into stomata (Robinson et al., 2011). SLGCs can immediately differentiate or generate new meristemoid/SLGC pairs through “spacing divisions” precisely oriented to avoid placing meristemoids next to existing stomata (Geisler et al., 2000). Polarity and cell-cell signaling regulate ACD propensity and orientation, resulting in stomata obeying a “one-cell spacing” rule (Lee and Bergmann, 2019). Loss of *BASL* has profound effects on division plane placement, self-renewing divisions, and cell fates in meristemoids and SLGCs, resulting ultimately in the accumulation of clustered stomatal precursors and clustered stomata (green and purple shading, respectively, in Figure 1B). A central role for BASL in polarity and ACDs is supported by genetic and protein interaction experiments showing that BASL is required for asymmetric localization and inheritance of signaling cascades linked to cell fates (Guo et al., 2021; Zhang et al., 2016a; Zhang *et al*., 2015) as well as other polarity proteins (Houbaert et al., 2018; Pillitteri et al., 2011; Rowe et al., 2019).

**Figure 1.**
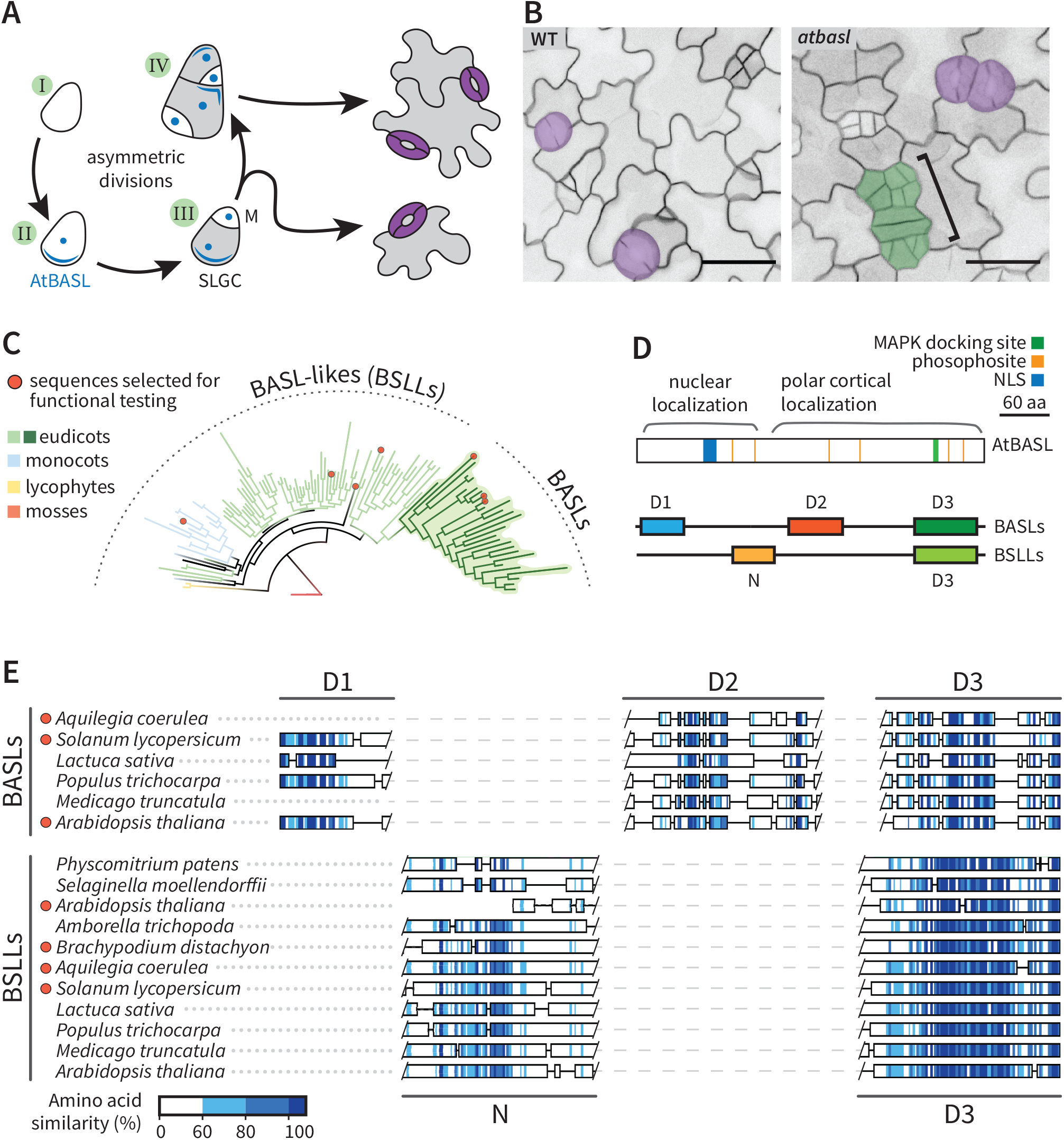
Stomatal asymmetric division regulator gene *BASL* homologues are restricted to eudicots. (A) Schematic of AtBASL activity during asymmetric cell divisions (ACDs) in the *Arabidopsis* stomatal lineage. BASL (blue) is expressed in progenitors entering the stomatal lineage (I), localizes the nucleus and to a polar crescent at the cell cortex before division (II) and is coordinated with the cell division plane to ensure its asymmetric inheritance (III). The larger, BASL-inheriting daughter is a stomatal lineage ground cell (SLGC) that may differentiate into a pavement cell (grey). The smaller, non-BASL-inheriting daughter is a meristemoid (M) that may differentiate into a stoma (purple). Both SLGCs and meristemoids, however, may undergo additional divisions requiring re-expression or reorientation of the BASL crescent (IV). (B) DIC images of the abaxial epidermis of WT and *atbasl* cotyledons, false colored with stomata in purple and clustered small cells in green. Scale bars: 30 µm. (C) Phylogenetic tree of protein sequences showing BASL clade arising from larger BSLL family. (D) Schematic of experimentally defined AtBASL domains and motifs from (Dong et al., 2009; Zhang et al., 2015) and major conserved domains in BASLs and BSLLs, shown approximately to scale. (E) Amino acid similarity scores (see Methods) displayed over two independent alignments of putative BASL and BSLL orthologues, annotated with major domains of sequence conservation in BASLs (D1 – D3) and BSLLs (N and D3). *See also Figure S1*.

Despite a pivotal role in Arabidopsis stomatal lineage development, and the fact that asymmetric divisions have been observed in the stomatal lineages of angiosperms, gymnosperms and ferns (Rudall et al., 2013), it is not known if BASL acts in diverse stomatal lineages. This is due, in part, to debates as to whether *BASL* homologues are even present in other plants (Chan et al., 2020; Vatén and Bergmann, 2012). Here we use a combination of phylogenetic analyses, cross-species complementation, and functional analysis in native contexts to identify likely *BASL* orthologues. We find proteins that can *p*olarize and function as BASL are restricted to the dicots but are related to more broadly distributed proteins, suggesting that BASL may be built from an ancestral MITOGEN-ACTIVATED PROTEIN KINASE (MAPK) interacting domain chassis, to which polarization capacity was added. Analysis of *SlBASL* dynamics and loss-of-function mutants in tomato reveal conservation in cell autonomous BASL roles, but unique implementation of those roles in the context of divergent patterning regimes, including novel cell fate transitions and spacing mechanisms. This multi-species analysis provides insight into the evolution of a unique polarity regulator and into the developmental choices available to cells as they build coordinated tissues and organs.

## RESULTS

### BASL is a rapidly evolving eudicot-specific gene

Previous efforts to reconstruct the evolutionary history of BASL have been hampered by low overall sequence conservation beyond close relatives of Arabidopsis (Chan *et al*., 2020; Vatén and Bergmann, 2012), although select protein domains could be found in more distantly related species (Zhang *et al*., 2016a). Using a reciprocal best hit heuristic, we found a rapid drop-off in sequence similarity of candidate BASL orthologues outside the Brassicaceae, with a plateau of around 60% amino acid similarity (30% identity) for eudicots followed by a sharp drop to ∼30% similarity (10% identity) for non-eudicot candidates (Figure S1A). To untangle the relationships between these sequences, we queried the UniProt KB database using jackhmmer and a high-quality reference alignment of putative BASL orthologues from the Brassicaceae (for sequences and trees used in this paper, see Supplemental Data on figshare).

This relatively permissive search strategy returned many hundreds of putative orthologues across the plant kingdom (Figure 1C). Inspection of hits from 33 land plants revealed that eudicot genomes typically contain a single BASL orthologue candidate with two or three short but well conserved domains in common with AtBASL (termed D1 – D3) (Figures 1D-E and S1B and Table S1). BASL candidates in this group also show conserved splice junctions, a shared profile of ordered and intrinsically disordered regions and generally concordant secondary structure predictions at predicted ordered sites (Figure S1C). A second group of sequences shows strong conservation surrounding a previously identified MAPK docking site (FxFP or DEF site, (Jacobs et al., 1999)) located in the D3 domain (Zhang *et al*., 2015) but are not otherwise similar to AtBASL. These BASL-like (BSLL) sequences also share a unique N-terminal domain (termed N) (Figure S1D-E and S1B). BSLLs are present in the genomes of both eudicot and non-eudicot land plants, including the distantly related moss *Physcomitrium patens*, but not the alga *Chlamydomonas reinhardtii* or the liverwort *Marchantia polymorpha*, which lack stomata. These data suggest that BASL may have arisen via duplication from an ancient clade of BSLL proteins encoding a MAPK docking site embedded in the D3 domain. In eudicots, addition of a structured domain at the D2 position may have created a distinct BASL clade, given that functional AtBASL requires both D2 and D3 domains (Dong *et al*., 2009). Some eudicot orthologues also contain an N-terminal D1 domain; this domain is dispensable for AtBASL function (Dong *et al*., 2009) and has been secondarily lost in several lineages, including legumes such as *Medicago truncatula*. (Figure 1E and S1C).

### BASLs, but not BSLLs, polarize in developing leaves

*BSLL* genes have not been characterized in any species and, despite the absence of D1 and D2 domains, our phylogenetic analyses do not preclude the possibility that BSLL proteins may be functionally redundant to BASL outside of Arabidopsis. In fact, *BSLL* gene expression is enriched in the stomatal precursor zone at the base of rice and maize leaves (Wang et al., 2014). Therefore, to characterize BASL and BSLL homologues, we began by assaying the feature that best defines BASL—its polarized subcellular localization, which can be replicated in non-native contexts (Chan *et al*., 2020; Dong *et al*., 2009; Zhang *et al*., 2015). We expressed BASL and/or BSLL coding sequences from *Arabidopsis thaliana, Solanum lycopersicum, Aquilegia coerulea* and the monocot outgroup *Brachypodium distachyon* in developing *Nicotiana benthamiana* leaves (Figure S2A-B). We found that proteins encoded by putative *BASL* orthologues from tomato and *Aquilegia* localized to a polar crescent at the edge of pavement cell lobes in *N. benthamiana*, much like AtBASL, although only AtBASL accumulated in the nucleus (Figure S2A). In contrast, all BSLL candidates exhibited diffuse cortical localization with no visible polarized domain (Figure S2B).

Based on these data, we selected three candidates for more extensive functional analysis: the tomato protein SlBASL (Solyc3g114770), which conserves all three major domains, but displays modest overall sequence conservation (23% AA identity and 50% AA similarity); the *Aquilegia* protein AqBASL (Aqcoe7g067300), which conserves only D2 and D3 domains; and the *Brachypodium* protein BdBSLL1 (Bradi3g47127), which is the protein most similar to AtBASL in that genome but whose similarity is restricted to the D3 domain (see Table S3 for gene names and gene codes used in this paper).

We expressed these candidates in Arabidopsis as fluorescently tagged fusion proteins under control of the native *AtBASL* promoter. Both SlBASL (AtBASLpro::VENUS-SlBASL) and AqBASL (AtBASLpro::VENUS-AqBASL) reporters exhibited robust polarization, with the polar domain consistently located in the larger daughter cell post-division, similar to a native AtBASL reporter (Figure 2A-2C). In contrast, expression of a BdBSLL1 reporter (AtBASLpro::VENUS-BdBSLL1) was cortically enriched but failed to polarize and was inherited symmetrically by both daughters following division (Figure 2D).

**Figure 2:**
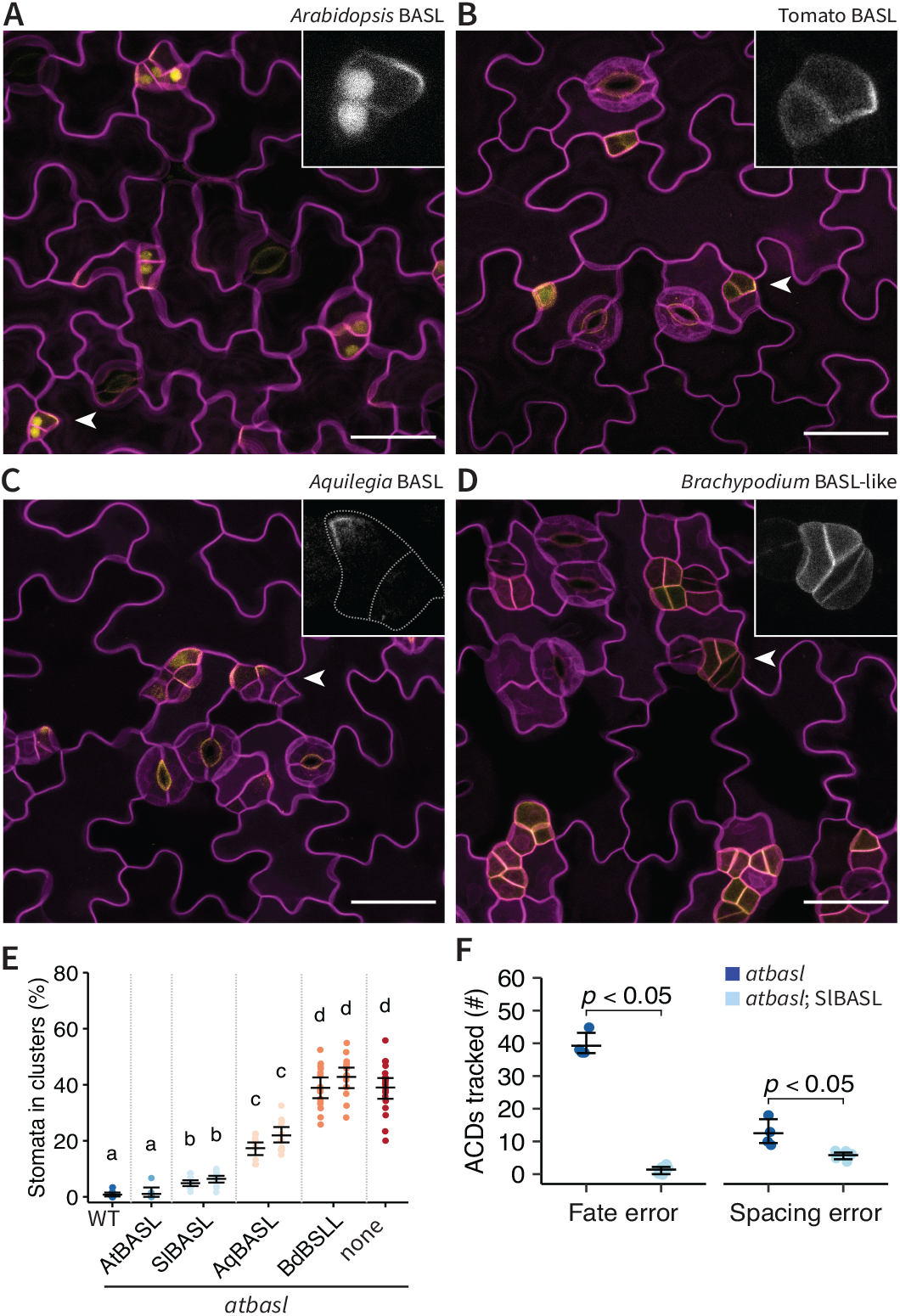
BASLs, but not BSLLs, polarize and rescue the *atbasl* mutant. (A-D) Confocal images of BASL and BSLL translational reporters (yellow) in abaxial epidermis of 4 dpg *atbasl* cotyledons. Cell outlines (magenta) visualized by *ML1p::mCherry-RCI2A*. White arrowheads mark cell enlarged in inset to highlight reporter distribution during ACD. Scale bars: 30 µm. Cells were hand-outlined in inset (C) for clarity. (E) Quantification of stomatal clustering phenotypes in *atbasl* as rescued by reporters noted on x-axis. For SlBASL, AqBASL and BdBSLL constructs, results from two independent lines are shown. n = 120 stomata in each of 8 – 20 leaves. For letters above each sample, see methods. (F) Quantification of the origin of stomatal pairs in *atbasl* and the *atbasl*; SlBASL rescue line. n = all trackable pairs in each of 4 leaves. Numerical data in E - F are represented as mean ± 95% confidence interval. Bonferroni-corrected p-values from Mann-Whitney *U* test are shown. *See also Figure S2*.

The terminal phenotype of *atbasl* mutants is clustered stomata, which can originate from two distinct errors: misoriented spacing divisions or failures to differentiate cell fates after ACD (Gong et al., 2021a). We measured the ability of BASL and BSLL reporter constructs to rescue the stomatal clustering phenotype of *atbasl* mutants and observed a graded response: while an AtBASL construct nearly perfectly rescue the mutant phenotype (99% rescue capacity, details in methods), rescue was partial with SlBASL (87%) and modest with AqBASL (50%) constructs (Figure 2E). As expected from its lack of polarization, BdBSLL1 had no significant capacity to correct stomatal clustering (Figure 2E). Finer dissection of SlBASL rescue function through lineage tracing showed that SlBASL nearly completely rescued stomatal pairs that arose from fate errors, but was only moderately effective at correcting spacing errors (Figure 2F).

Together, these data suggest that among dicots, there are true BASL orthologues that retain polarization capacity and the ability to regulate division plane orientation and fate asymmetry during ACDs. BSLLs, as exemplified by BdBSLL1, do not. The apparent lack of functional BASL orthologues in the grasses may be related to the differences in ontogeny of grass stomata, where stomatal guard cell formation is preceded by a single ACD that is invariantly oriented relative to the overall leaf axis (McKown and Bergmann, 2020). Interestingly, we found that an AtBASL reporter expressed in leaves of the pooid grass *Brachypodium distachyon* polarly localized in crescents oriented toward the base of the leaf and was preferentially expressed in the larger daughter cells resulting from ACDs (Figure S2D). In newly formed stomatal complexes, AtBASL also localized to the junction between guard cells and subsidiary cells (Figure S2D). These behaviors indicate that BASL can recognize tissue-wide polarity information created through other, more ancient, polarity networks, even if it is not a functional component of those networks itself.

### SlBASL is polarized and involved in stomatal pattern in tomato

If proteins sharing AtBASL’s polar localization exist outside of Brassicaceae, and can substitute for AtBASL in ACDs of the Arabidopsis stomatal lineage, what is their role in their native species? To address this, we generated reporters and CRISPR/Cas9-induced mutation lines to test localization and function of BASL in another dicot. We chose *Solanum lycopersicum* (tomato) for these analyses because of its phylogenetic position among dicots, efficient transformation protocols and the rich history of developmental and physiological studies on tomato leaves and stomata (Goliber et al., 1998; Ichihashi et al., 2014; Nir et al., 2017).

Stomata on the M82 tomato cotyledon and leaf are spaced to avoid direct contact and mature organs feature stomata distributed among lobed pavement cells in a pattern resembling that found in mature Arabidopsis leaves (Figure 3D and S3A). Earlier, during cotyledon development (3 days post germination (dpg) on MS-agar plates), when small epidermal cells with the morphological characteristics of meristemoids are abundant, we begin to our SlBASL translational reporter (SlBASLpro::VENUS-SlBASL) (Figure 3A). In still images, SlBASL appeared in a polarized cortical crescent consistently associated with the larger daughter cell after ACD (inset in Figure 3A) but lacked the symmetrically inherited nuclear localization typical of AtBASL (Figures 2B and 3A). To monitor SlBASLpro::VENUS-SlBASL dynamics during a single cell division cycle, we used time-lapse imaging, collecting images at 30 min intervals from 1 dpg abaxial cotyledons. As shown for two representative cells in Figure 3B (n = 19 cells tracked), SlBASL first appeared several hours before division but was diffuse and depolarized. SlBASL expression remained quite weak during cell division but intensified and became distinctly polarized post-division (Figure 3B, white arrowheads). The onset of SlBASL polarization differs from AtBASL, which is typically polarly localized by the beginning of mitosis in time to direct orientation of the division plane (Gong et al., 2021c).

**Figure 3.**
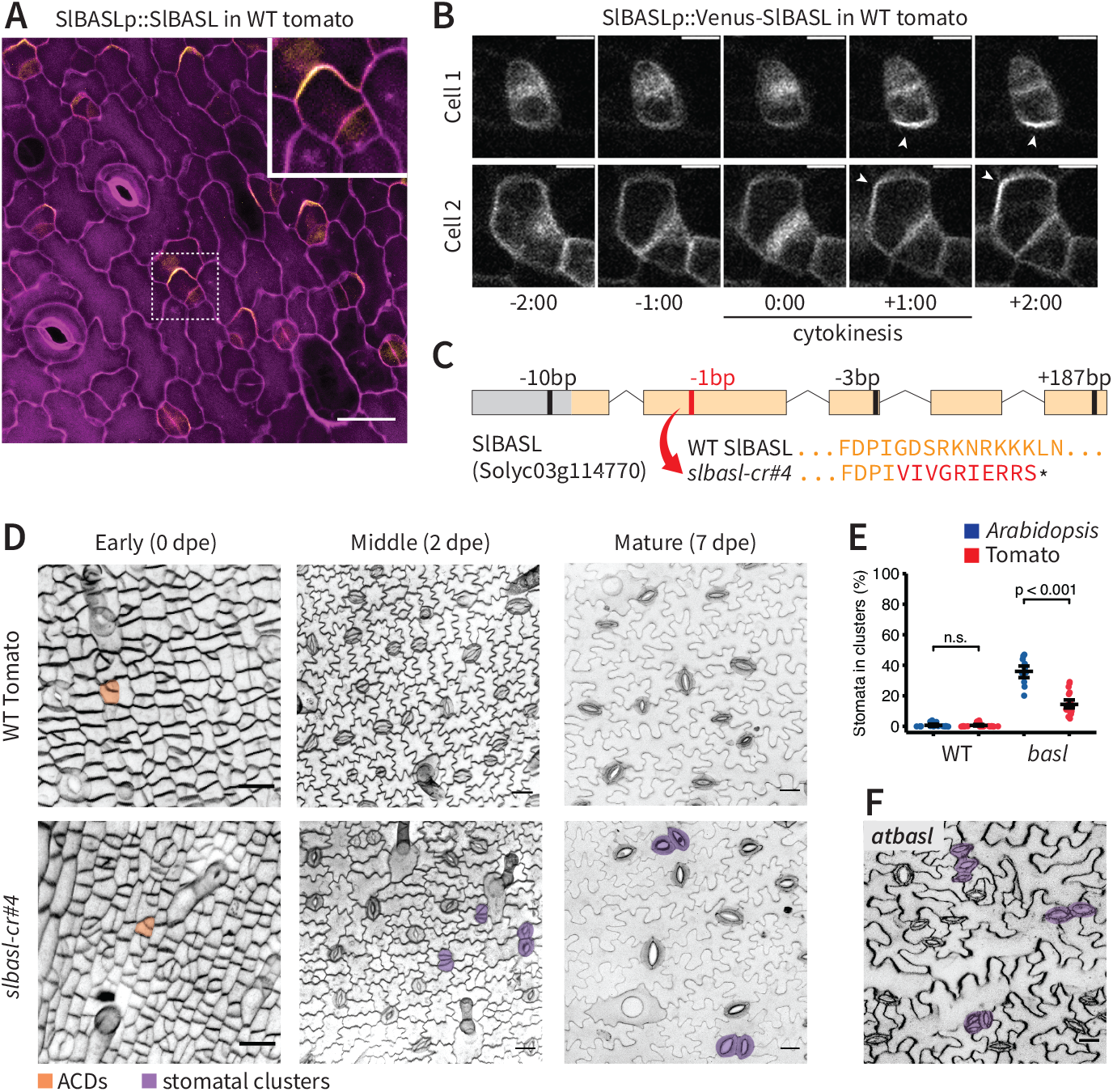
SlBASL is polarized and required for proper stomatal patterning in tomato. (A) Confocal image of *SlBASLp::Venus-SlBASL* (yellow) expression in abaxial epidermis of 1 dpe cotyledon. Cell outlines (magenta) visualized by propidium iodide (PI) staining. Dotted line marks cells shown in inset. Scale bar: 30 *µ*m. (B) Time-lapse of SlBASL reporter localization in two representative cells undergoing ACDs. Arrowheads indicate polarized SlBASL. Time shown as hours:minutes. Scale bar: 10 *µ*m (C) Schematic of *slbasl-cr#4* allele. Black and red bars indicate the positions of gRNAs along the transcript. In red, a 1 base pair deletion leads to a premature stop codon after 63 amino acids. Wildtype and mutant protein sequences surrounding this frameshift are shown below. (D) Confocal images of the abaxial epidermis of M82 and *slbasl-cr#4* cotyledons at indicated ages. Cell outlines visualized by FM4-64 staining. Example ACDs are false colored in orange, stomatal pairs in purple. Scale bar: 30 *µ*m. (E) Quantification of stomatal clustering in the abaxial epidermis of wild type and *basl* mutants of *Arabidopsis* (14 dpg) and tomato (10 dpe). Clusters are reported as the fraction of stomata in direct contact with other stomata/total stomata in mature cotyledons Data are represented as mean ± 95% confidence interval. Bonferroni-corrected p-values from Mann-Whitney *U* test are shown. n.s., not significant. (F) Confocal image of stomatal clusters (purple) in *atbasl* mutant epidermis (14 dpg). *See also Figure S3*.

Given SlBASl’s polar localization and its ability to complement *atbasl*, we hypothesized that it would be required for stomatal patterning in tomato. We targeted the *SlBASL* locus for CRISPR/Cas9 mutagenesis using multiplexed guides at the 5’UTR, and exons 2, 3 and 5 and obtained three independent lines bearing mutations at the *SlBASL* locus (*slbasl cr#2, cr#4, cr#6*, Figures 3C and S3E). In the T2 generation, plants bearing these *slbasl* mutations were not noticeably different from control M82 plants in overall size, growth habit or fertility, but each line displayed stomatal clusters on the epidermis, suggesting that SlBASL function was disrupted (Figure 3D and S3B). For detailed phenotypic and quantitative characterization, we selected the *slbasl-cr#4* allele, where a 1 base pair deletion in exon 2 is predicted to cause a frameshift and premature stop codon after 63 amino acids (shortly after domain D1, Figure 3C and S3E-F).

We imaged wild type (M82) and *slbasl-cr#4* soil-grown cotyledons representing early (0 days post emergence, dpe), young (2 dpe) and mature (7 dpe) stages of epidermal development. At early stages, many small box-like cells and physically asymmetric divisions (Figure 3D. orange shading) could be observed in both M82 and *slbasl-cr#4*. At 2dpe, lobed pavement cells and all classes of stomatal lineage cells—meristemoids, SLGCs, GMCs and mature guard cells—could be identified in both genotypes, but *slbasl-cr#4* also exhibited occasional stomatal pairs (Figure 3D, purple shading). At maturity (7 dpe), the abaxial cotyledon and true leaf epidermis of *slbasl-cr#4* plants exhibited many clustered stomata relative to wild type plants (Figure 3D and S3B).

Although loss of *SlBASL* resulted in stomatal clusters in all scored cotyledons (Figure 3D), the phenotype in tomato was much milder than that generated by loss of *AtBASL* in Arabidopsis (Figure 3E).

Because we had identified multiple mutant alleles of *SlBASL* (Figure S3B) and because we found no evidence for sequences that could encode redundant *SlBASL* genes (see Methods), we hypothesized that differences in severity of *basl* phenotypes were not due to incomplete loss of *BASL* activity, but rather might reflect different strategies for stomatal spacing between these two species.

To track stomatal development over time, and to identify the origin of the stomatal pairs in *slbasl*, we used a long-term lineage tracing strategy similar to (Gong *et al*., 2021a) where daughters of each ACD were following through subsequent divisions and differentiation to terminal cell fates. We adapted this time-course and lineage tracing method for tomato by creating a live-cell marker for epidermal cell outlines (ML1p::RCI2A-mNeonGreen), and following epidermal development in soil-grown plants starting at 1 dpe to capture the widest variety of division competent cell types (e.g. Figure 4A-4B, orange).

**Figure 4.**
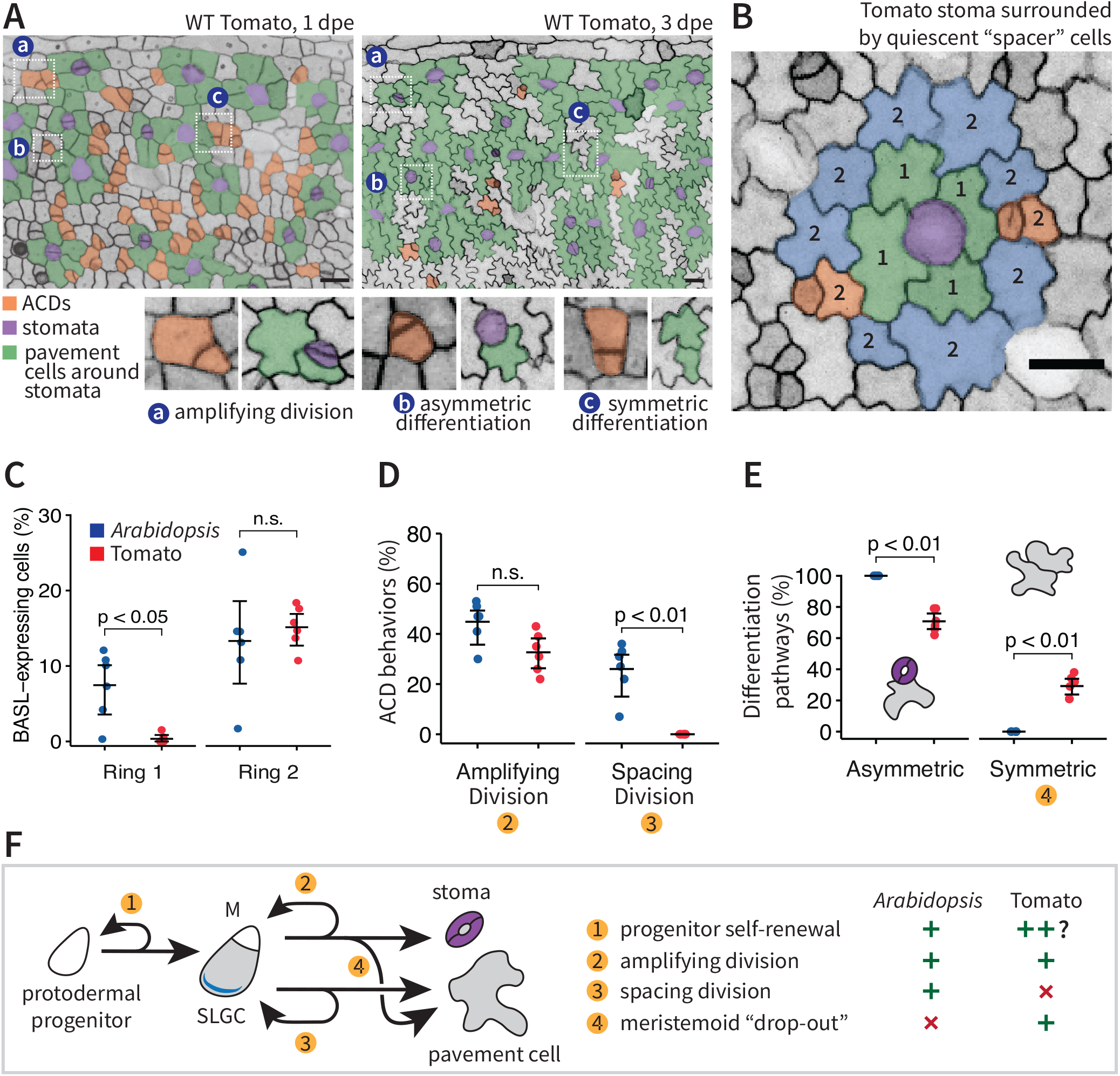
Rewired fate transitions in tomato stomatal development. (A) Confocal images of developmental time-course of abaxial cotyledon epidermis. Cell outlines visualized by *ML1p::RCI2A-NeonGreen*. White boxes mark division and differentiation events (a-c) shown below. Cell types are false colored as indicated in key. Scale bars: 50 *µ*m. (B) Image to illustrate typical spatial organization of cells surrounding stomata (purple) at 4dpe, including cells immediately adjacent (ring 1, green) and one cell removed (ring 2, blue and orange). Scale bar: 30 *µ*m. (C) Percentage of cells expressing BASL in successive rings surrounding stomata. n = ∼250 cells in each of 6 leaves. (D) Percentage of ACDs that underwent additional amplifying or spacing divisions. ACDs displaying both amplifying and spacing divisions in our imaging window (2 days) are counted once for each class of division. n = ∼45 ACDs in each of 6 imaging fields. (E) Percentage of ACDs that resolve into asymmetric (one stomata and one pavement cell) or symmetric (two pavement cells) cell fates. n = ∼45 ACDs in each of 6 imaging fields (∼0.125 mm^2^ at 1dpe; ∼0.3 mm^2^ at 3 dpe) (F) Model of symmetric and asymmetric division types that contribute to stomatal fate and pattern featured in *Arabidopsis* and tomato stomatal lineages, with shifts in predominant types between the species summarized to the right. Numerical data in C – E are represented as mean ± 95% confidence interval. Bonferroni-corrected p-values from Mann-Whitney *U* test are shown. n.s., not significant.

When cells were tracked between 1 dpe and 3 dpe timepoints, we readily observed symmetric GMC divisions and asymmetric divisions of meristemoids (amplifying divisions, Figure 4A). However, asymmetric divisions of SLGCs (spacing divisions), were nearly undetectable (Figure 4D). As misoriented spacing divisions are major contributors to pattern defects in *atbasl* mutants, this immediately suggested a reason for the mild *slbasl* phenotype in tomato. Indeed, seven out of eight lineage-tracked stomatal pairs in *slbasl-cr#4* arose from a fate defect (Figure S3C-D).

If spacing divisions are not major contributors to epidermal pattern in tomato, then other patterning mechanisms may enforce 1-cell spacing. We therefore returned to the time course images, focusing on outcomes of asymmetric divisions and on the behaviors of cells surrounding young stomata. We found that spacing (and all other) divisions were suppressed in the first ring of cells around a stoma (Figure 4B-C), as was SlBASL expression (Figure 4C). In contrast, AtBASL expression in *Arabidopsis*, where spacing divisions are common, was not excluded from ring 1 (Figure 4C). Because the *SlBASL* promoter can drive expression during spacing divisions in transgenic *Arabidopsis* plants (Figure S2D) and conserves several SPCH binding motifs that may drive expression during ACDs (Figure S2E), the exclusion of SlBASL from this first ring of cells likely results from changes to the developmental system upstream of *SlBASL*. Interestingly, although asymmetric spacing divisions were reduced, there were still many physically asymmetric divisions in the tomato epidermis (Figure 4A, orange). Unexpectedly, many resolved symmetrically as pairs of pavement cells (Figure 4A and 4E), a phenomenon never observed in wild type Arabidopsis. We termed this novel fate transition “meristemoid drop-out” due to the loss of stomatal precursors from the lineage. Considered together, the relationships between morphologically asymmetric divisions and the resultant daughter cell fates and behaviors point to remarkably divergent developmental trajectories converging on similar final epidermal patterns in Arabidopsis and tomato (Figure 4F).

## DISCUSSION

Our data suggest that BASL is a recently evolved eudicot-specific plant polarity protein that has been coopted into more ancient programs of asymmetric divisions in stomatal development. Comparison of BASL expression and function in Arabidopsis and tomato, however, revealed that even among eudicots, the way polarity and ACDs are employed during stomatal development can be quite different (Figure 4F). In tomato cotyledons there is a near-complete lack of spacing divisions and divisions surrounding young and mature stomata, but in Arabidopsis, ACDs adjacent to stomata are common and their occurrence and orientation must be carefully regulated to avoid stomatal clustering. This regulation is disrupted in the *atbasl* mutant, where ∼30% of stomatal clusters arise through defects in division orientation (Gong *et al*., 2021a). In contrast, stomatal development in tomato employs quiescence of both sister and non-sister cells surrounding stomata to create a pavement cell “buffer zone” (Figure 4B) that ensures a degree of stomatal spacing even in the *slbasl* mutant. Additionally, in tomato, approximately 30% of physically asymmetric divisions undergo “meristemoid drop-out” and resolve as pairs of pavement cells (Figure 4E). While such divisions are absent in wild type Arabidopsis development, similar phenotypes have been described in *myoxi-i* mutants, where nuclear migration and division plane defects accompany differentiation of both daughters into pavement cells (Muroyama et al., 2020). Similarly, late depletion of the transcription factor SPEECHLESS can “divert” meristemoids or GMCs towards pavement cell fate (Lopez-Anido et al., 2021). In tomato, such decoupling of physical and fate asymmetry can be used to populate the developing epidermis with additional pavement cells and promote proper spacing of stomata.

Different strategies for spacing stomata apart may incur different costs. As spacing divisions allow Arabidopsis to generate additional meristemoids under favorable environmental conditions (Gong et al., 2021b; Vatén et al., 2018), their lack in tomato may represent a constraint on the plasticity of stomatal development. Plasticity could be restored, however, by regulating flux along other paths in the lineage, such as in the number of entry divisions, or by converting “drop-outs” into meristemoids (Figure 4F). Alternatively, a latent spacing division capacity may be activated under appropriate environmental conditions not tested in our assays, as the SlBASL promoter appears to contain the necessary *cis*-regulatory information to operate in spacing divisions when expressed in Arabidopsis.

The paucity of spacing divisions in tomato may have relaxed selection on SlBASL protein activity, an idea supported by its effective rescue of *atbasl* stomatal pairs originating from fate defects, but only modest rescue of pairs originating from spacing divisions (Figure 2F). When considering sources of specificity, it is notable that although highly diverged, BASL orthologues from eudicots conserve three distinct protein domains, including a previously described MAPK docking site at the C-terminus (Zhang *et al*., 2016a) and two newly identified domains at D1 and D2 positions. Consistent with previous dissections of AtBASL domain structure (Dong *et al*., 2009), BASL orthologues conserving only D2 and D3 domains (as exemplified by AqBASL) appear to be sufficient for substantial rescue of the *atbasl* phenotype, despite broad divergence elsewhere in the peptide sequence.

Among these domains, D3 is most highly conserved and present in BSLLs, an ancient clade of proteins that do not polarize. This domain overlaps several residues where MAPKs phosphorylate AtBASL (Zhang *et al*., 2015) and a conserved specialized MAPK docking motif (FxFP) that, although phylogenetically widespread, occurs in only a modest number of proteins in each species (Fernandes and Allbritton, 2009). We speculate that D3-containing BSLL genes in early diverging angiosperms may have provided a MAPK-interacting “chassis” to which a polarity module was linked in eudicots, and the resultant protein became useful to enforce cell fate segregation during stomatal lineage ACDs.

BASL’s ability to polarize also requires information encoded in the D2 domain. This domain overlaps a region of AtBASL shown to interact with BREVIS RADIX (BRX) family proteins (Rowe *et al*., 2019), which exhibit mutual dependency with BASL for polarization and function in the Arabidopsis stomatal lineage. In turn, BRX interacts with BASL through a pair of conserved BRX domains (Rowe *et al*., 2019), which are also found in other proteins such as PRAFs (Briggs et al., 2006). Interestingly, BASL may interact with the BRX domain through a pair of well-conserved β-strands in the D2 domain, which bear similarity in structure, but not sequence, to the BRX-interacting domain of LAZY3, a known PRAF binding partner (Furutani et al., 2020). These data suggest possible convergence of a BRX interaction fold across two plant-specific polar protein families: BASLs and LAZYs. Convergent evolution of protein-protein interaction folds among polar proteins has also been described for DIX domains, which are found in animal Disheveled and plant SOSEKI proteins (van Dop et al., 2020).

Regardless of how BASL homologues are refined for participation in different types of ACDs, we are still left with the conundrum of how non-eudicots coordinate ACDs without BASL. Stomatal development has been extensively studied in grasses such as maize, rice and *Brachypodium distachyon*, where meristemoids typically undergo a single, uniformly oriented asymmetric division before terminal differentiation (McKown and Bergmann, 2020). This mode contrasts with the flexible program exhibited by many eudicots, including *Arabidopsis* and tomato, where the number and orientation of asymmetric divisions is variable and responsive to a variety of internal and external inputs. It is tempting to speculate that BASL may have been coopted in eudicots to coordinate some aspect of this flexible developmental program that is dispensable in grasses. Alternatively, BASL functions in non-eudicots may be performed by an unrelated protein family, such as BRX family proteins, which polarize alongside BASL in the *Arabidopsis* stomatal lineage and appear to be deeply conserved across land plants (Briggs *et al*., 2006; Ramalho et al., 2021; Rowe *et al*., 2019). Recent work (Ramalho *et al*., 2021) to reconstruct the long-term evolutionary history of *Arabidopsis* proteins, combined with studies of protein function in their native contexts, may shed light on the genetic toolkit and behavior potential available to ancient and modern stem cells as they progress through development.

## MATERIALS AND METHODS

### RESOURCE AVAILABILITY

#### Lead Contact

Further information and requests for resources should be directed to and will be fulfilled by the Lead Contact, Dominique Bergmann.

#### Materials Availability

There are no restrictions on materials generated for this manuscript.

#### Data and Code Availability

Multiple sequence alignments and Newich tree files are available on figshare (doi:10.6084/m9.figshare.15109575)

### EXPERIMENTAL MODEL AND SUBJECT DETAILS

*Arabidopsis thaliana* Col-0 seeds were surface-sterilized with 75% ethanol and stratified for 2 days. After stratification, seedlings were grown on ½ strength Murashige and Skoog (MS) media (Casson Labs) with 1% agar for 3 – 14 days under long-day conditions (16 hr light/8 hr dark at 110 μmol m^2^ s^-1^ and 22°C).

*Brachypodium distachyon* Bd21-3 seeds were stratified for 2 – 8 days. After stratification, seedlings were grown on agar-solidified ½ MS media at 22-26 °C, at 80 μmol m^2^ s^-1^ and 16 hr light/8 hr dark cycles. For propagation, plants were transferred to soil in a greenhouse (20 hr light/4 hr dark, 250-300 μmol m^2^ s^-1^; day temperature: 28°C; night temperature: 18°C).

Tomato plants (*Solanum lycopersicum* cv. M82) were grown in a growth room set to a photoperiod of 16 hr light/8 hr dark, light intensity of approximately 250 µmol m^2^ s^-1^, and 26 °C. For propagation, plants were grown in a greenhouse under natural day length conditions, at 700–1200 µmol m^2^ s^-1^ and 18 – 29 °C.

## METHOD DETAILS

### Identification of BASL and BSLL sequences

BASL sequences from 9 Brassicaceae species were aligned with MUSCLE (Edgar, 2004) and used to query the Viridiplantae section of the UniProtKB protein sequence database with jackhmmer (Finn et al., 2011; Potter et al., 2018). Candidate homologs from 33 land plant species with e-values below 0.01 were selected for alignment and manual inspection. After removing exact and near duplicates (>95% identity), sequences were aligned using MAFFT (Katoh and Standley, 2013) and sites with >80% coverage were used to estimate the phylogeny with the neighbor-joining algorithm as implemented on Geneious Prime 2.3. The tree shown in Fig. 1C is trimmed to remove several clades of highly diverged, likely spurious orthologues.

In several cases, BASL orthologues appeared to be missing in eudicot proteomes as annotated on UniProtKB but could always be found in other proteome or genome annotations (Tables S1-2). Similarly, parts of conserved domains at the extreme N- and C-termini that appeared to be missing in protein models could often be found in-frame in the corresponding DNA sequence; protein sequences were manually corrected to include such domains in Figure S1C. For that figure, intrinsic disorder predictions were derived from DISOPRED3 (Jones and Cozzetto, 2015). Secondary structure predictions were derived from PSIPRED 4.0 (Jones, 1999) and the confidence score of the prediction is used to color the alignment. The location of splice junctions was derived from the primary transcript annotated on Phytozome v12 (Goodstein et al., 2012) and exons were then shaded in alternating colors.

### Identification of BASL pseudogenes in tomato

A single BASL orthologue and three clearly distinguishable BSLL genes were found in the *Solanum lycopersicum* iTAG2.4 proteome. In the iTAG2.4 genome, one additional short ORF located within an intron of Solyc01g009450.2.1 could be recognized. Sequence inspection showed remnants of conserved D1 and D2 domains, suggesting orthology to SlBASL; however, several early stop codons and a frameshifting indel are predicted to produce a severely truncated 20 amino acid polypeptide from this locus.

### Arabidopsis and tobacco DNA constructs and plant transformation

BSLL candidates from tomato, *Aquilegia coerulea* and *Brachypodium distachyon* and a BASL candidates from *Aquilegia coerulea* and *Brachypodium distachyon* were synthesized *in vitro* following the primary annotated CDS from Phytozome v12. For transient expression in tobacco, constructs were cloned into the pH35YG or pICH47742 expression plasmids, introduced into *Agrobacterium tumefaciens* strain GV3101 and infiltrated into mature tobacco leaves using standard methods. For stable expression in *Arabidopsis*, constructs were cloned into the binary vector pAGM4723, introduced into *Agrobacterium* as above and transferred to *Arabidopsis* by floral dipping (Clough and Bent, 1998). Transgenic founder plants were identified by kanamycin resistance. All analyses were performed on homozygous T3 plants.

### Tomato DNA constructs and plant transformation

For tomato BASL marker lines, the BASL promoter and CDS were cloned into a level 0 MoClo part using the Golden Gate cloning system (Weber et al., 2011) and then fused to Venus and a NOS terminator to form a level 1 construct. The level 1 was transferred to a level 2, together with a kanamycin resistance cassette. For primers used in cloning see table below, red indicates Golden Gate tails. The constructs were sub-cloned into the pAGM4723 binary vector and were introduced into *Agrobacterium tumefaciens* strain GV3101 by electroporation. The constructs were transferred to M82 cotyledons, using transformation and regeneration methods described by (McCormick, 1991). Kanamycin-resistant T0 plants were grown and at least four independent transgenic lines were selected and self-pollinated to generate homozygous transgenic lines.

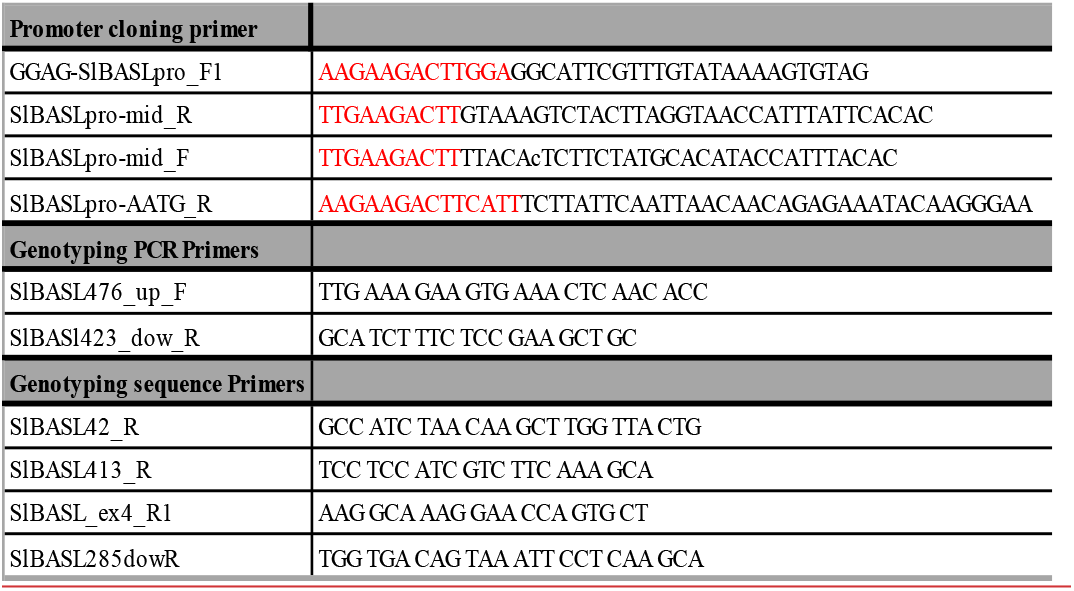

### *Brachypodium* experiments

For expression of AtBASL in *Brachypodium distachyon* strain Bd21-3, a Gateway pENTR plasmid encoding a fusion of YFP with the coding region of BASL (Dong *et al*., 2009) was recombined into the monocot transformation pIPKb002 that drives expression of inserts with the maize Ubiquitin promoter (Himmelbach et al., 2007) following standard Gateway protocols (Bragg et al., 2015). *Brachypodium* calli were transformed with AGL1 *Agrobacterium tumefaciens*, selected and regenerated according to standard protocols. Expression of the transgene was monitored in the development zone at the base of leaves from 22 dpg T1 plants as described in (Abrash et al., 2018).

### Tomato BASL CRISPR/Cas9 mutagenesis, plant transformation, and selection of mutant alleles

Four single-guide RNAs (sgRNAs) were designed using the CRISPR-P tool (Lei et al., 2014). The gRNAs and promoter were assembled using the Golden Gate cloning system as described in (Weber *et al*., 2011). The final binary vector including zCas9, the gRNAs and NPTII, assembled in pAGM4723, was introduced into *Agrobacterium tumefaciens* strain GV3101 by electroporation. The construct was transferred into M82 cotyledons using transformation and regeneration methods described by (McCormick, 1991). T0 transgenic plants resistant to Kanamycin were grown and independent lines were selected and self-pollinated to generate homozygous lines. For genotyping of the transgenic lines, genomic DNA was extracted, and each plant was genotyped by PCR for the presence of the zCas9. The positive lines for the zCas9 were further genotyped for mutations in SlBASL (Solyc03g114770) using a forward primer 400bp upstream to the ATG and a reverse primer 200 bp downstream to the stop codon, these pair primers cover the 4 gRNAs.

### Microscopy, image analysis and processing

All fluorescence imaging experiments on *Arabidopsis* plants were performed on a Leica SP5 confocal microscope with HyD detectors using 40x NA1.1 water objective with image size 1024*1024 and digital zoom from 1x to 2x. To quantify rescue of the *atbasl* stomatal clustering phenotype, 14 dpg cotyledons were imaged and the fraction of stomata in clusters of size two or larger was computed from regions containing 120 stomata (approximately 0.5 – 1.5 mm^2^ in size). Normalized rescue capacity was calculated as 1 – (% pairs in rescue - % pairs in WT) / (% pairs in *atbasl* - % pairs in WT). Still or time-course images of SlBASLpro::VENUS-SlBASL, ML1p::RCI2A-NeonGreen, propidium iodide (PI), and FM4-64 fluorescence in tomato were obtained from a Leica SP8 confocal microscope with HyD detectors using 20x oil objective with image size 1024*1024 and digital zoom from 1x to 2x. For time-course experiments on tomato, cotyledons of 2 dpe soil-grown seedlings were mounted with abaxial side toward coverslip, and 0.15% agarose was added to prevent root drying. Seedlings were carefully unmounted after imaging and returned to the soil until the next image acquisition. Time-lapse experiments of *SlBASL* in tomato were performed on a Leica SP5 confocal microscope with HyD detectors using 25x NA0.95 and 40x NA1.1 water objectives. Seedlings were mounted on a customized flow-through chamber (Davies and Bergmann, 2014) and images were acquired at 30 min. All raw fluorescence image Z-stacks were projected with SUM (*Arabidopsis* stills) or STD (tomato stills and time-lapse) Slices in FIJI. For all time-lapse images, drift was corrected after projection using the Correct 3D Drift plugin (Parslow et al., 2014) prior to any further analysis.

## QUANTIFICATION AND STATISTICAL ANALYSES

All statistical analyses in this manuscript were performed in RStudio (Team, 2020). Unpaired Mann-Whitney *U* tests were conducted with the wilcox_test function from the rstatix package (Kassambara, 2021). Bonferroni corrections were performed when more than 2 pair-wise comparisons were conducted, and Bonferroni-corrected p-values are indicated in all figures where applicable. For Figure 2E, each letter indicates a group of samples with no statistically significant difference between their means (p > 0.05).

