## Supplemental Figures 1-3 and Table S1-3 for "On the edge: Evolution of polarity protein BASL and the capacity for stomatal lineage asymmetric divisions"


Figure S1. Additional sequence features of BASL and BSLL proteins
Figure S2. Expression of BASL and BSLL proteins across species.
Figure S3. Additional phenotypes and molecular characterization of CRISPR/Cas9-generated *SlBASL* mutants


**Supplemental Tables**


Table S1: Number of BASL and BSLL proteins displayed in Fig. 1C
Table S2: Sequences used in Fig. S1C
Table S3: Gene names used in this paper


**Additional materials** hosted on figshare are available at doi:10.6084/m9.figshare.15109575.

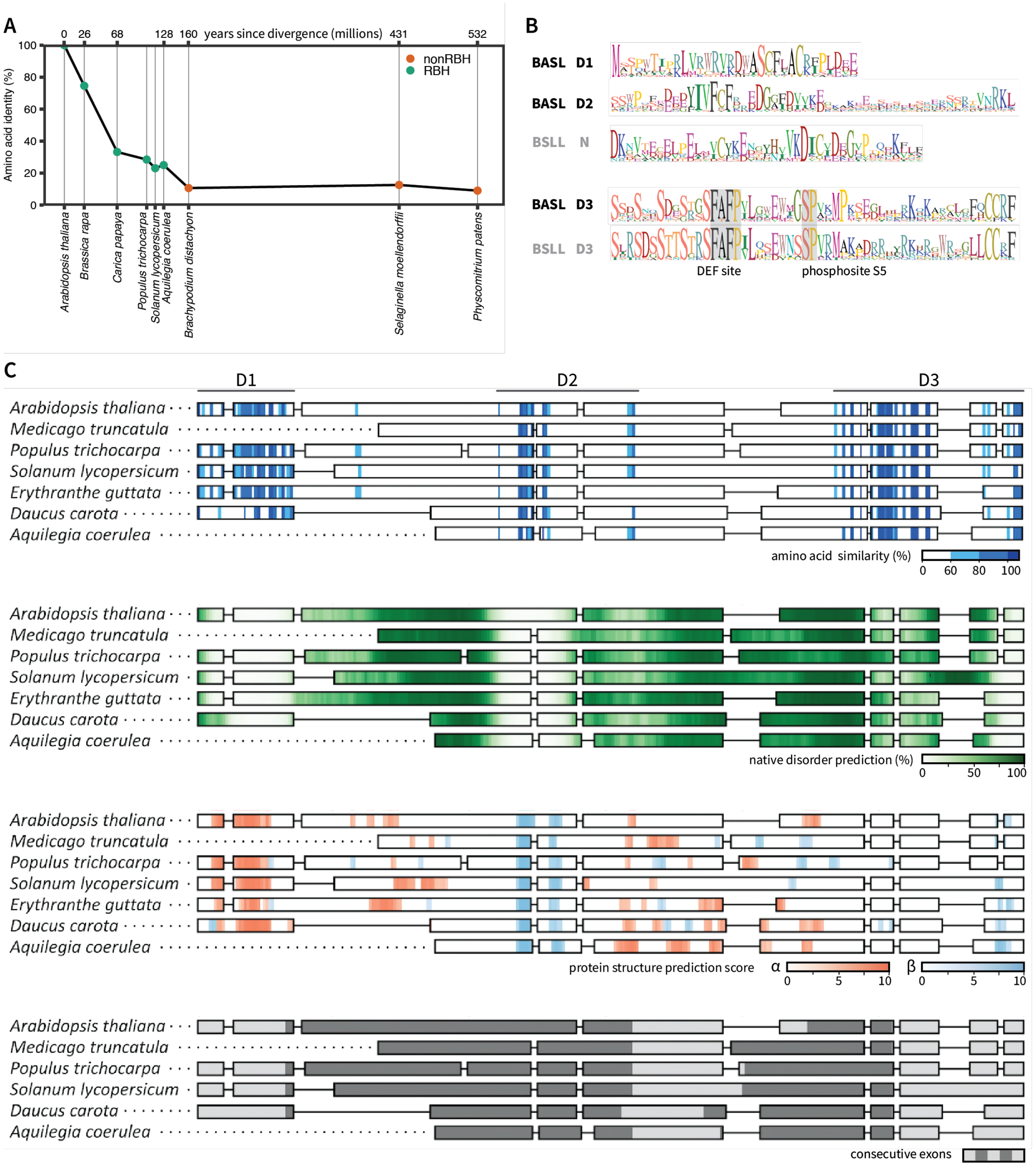


**Figure S1. Additional sequence features of BASL and BSLL proteins**

(A) Amino acid identity of top BLAST hits for AtBASL in select genomes. Sequences in green represent reciprocal best hits (RBH) when used to query the *Arabidopsis* genome. Note that all RBHs are from eudicot genomes, and eudicot RBHs show rapid decay in sequence identity outside the Brassicaceae (e.g., *Carica papaya*).

(B) Protein sequence logos of BASL and BSLL domains. In D3, grey boxes highlight a MAPK docking site (DEF) and phosphosite (S5) functionally characterized in *Arabidopsis* BASL (Zhang *et al*., 2016a; Zhang *et al*., 2015).

(C) Alignment of BASLs overlayed with amino acid similarity, predicted native disorder, predicted secondary structure and position of intron boundaries. Colors as indicated. In protein structure predictions, two PSIPRED scores are overlayed: α stands for α-helix, β for β-sheet.

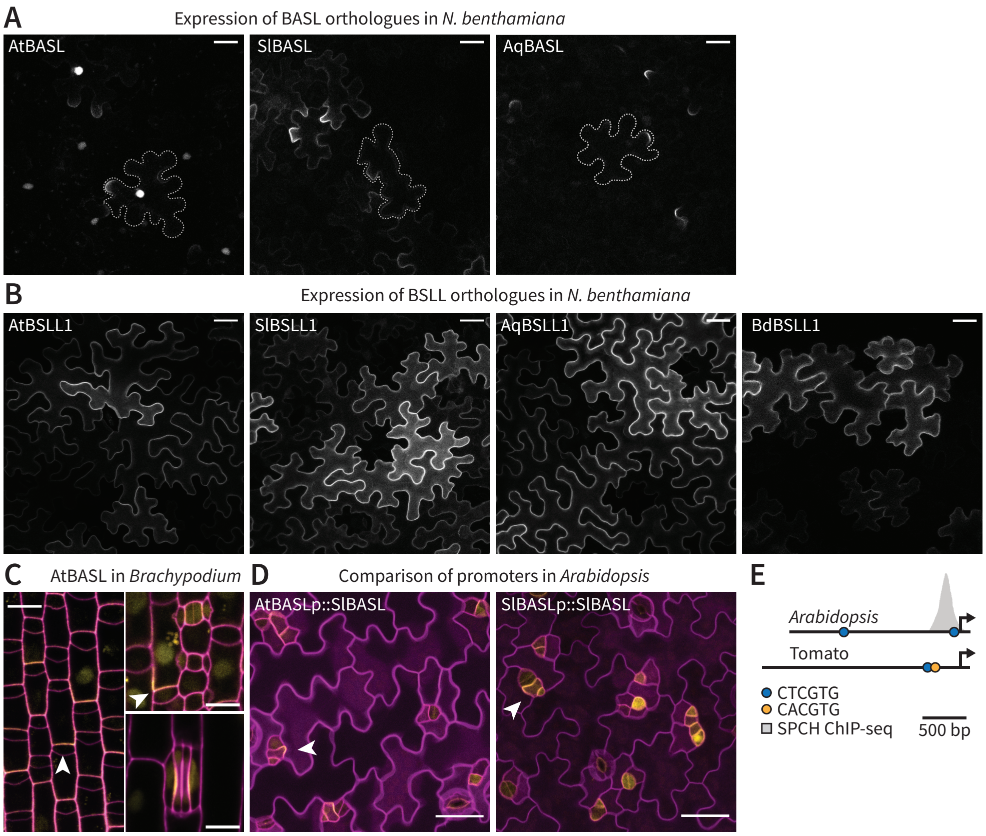


**Figure S2. Expression of BASL and BSLL proteins across species.**

(A) Confocal images of BASL candidate reporters polarly localized when transiently expressed in mature *N. benthamiana* leaves. Dotted outlines mark one representative cell. Scale bars: 40 $\mu$m.

(B) Confocal images of BSLL reporters when transiently expressed in mature *N. benthamiana* leaves. Scale bars: 40 $\mu$m.

(C) Confocal images of ZmUb::YFP-AtBASL (yellow) in *Brachypodium distachyon* leaves. Cell outlines (magenta) visualized by PI staining. Clockwise from left: expression during generative ACDs, in subsidiary cell progenitors and in a mature stomatal complex. White arrows indicate polarized localization. Images are oriented with leaf base towards the bottom. Scale bars: 10 $\mu$m

(D) Confocal images of SlBASL translational reporter (yellow) under the control of *Arabidopsis* or tomato *BASL* promoters in Arabidopsis cotyledon epidermis (cell outlines in magenta). Arrowheads mark polarized SlBASL accumulation during a spacing division.

(E) Schematic showing location of SPCH binding motifs (blue and yellow) in the promoters used in (D). AtSPCH ChIP-seq results from (Lau et al., 2014) are overlayed on the *Arabidopsis* promoter, showing binding near the proximal element

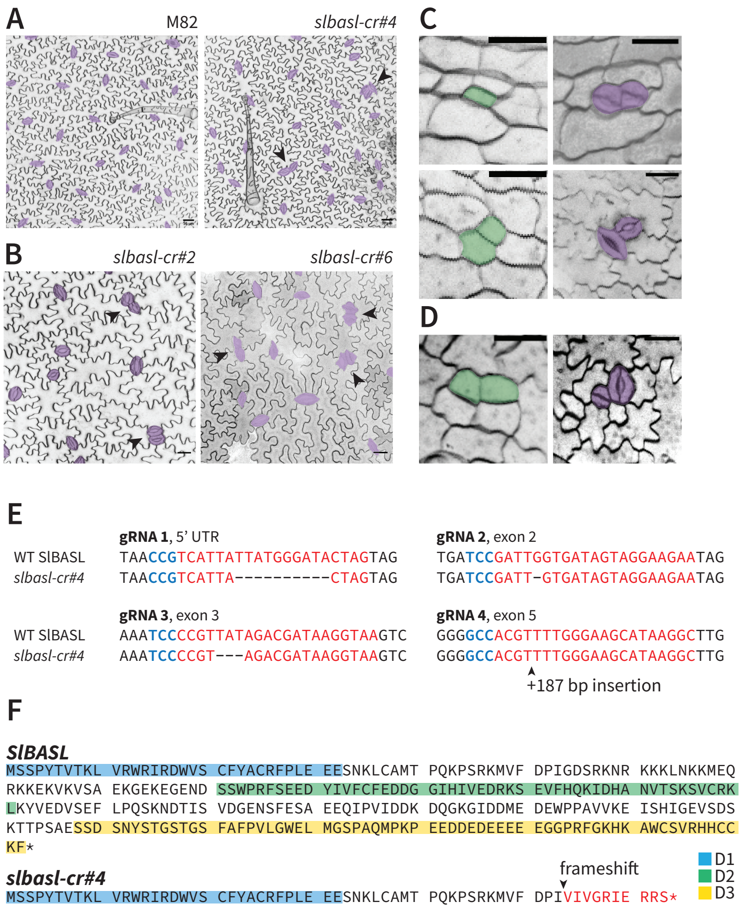


**Figure S3. Additional phenotypes and molecular characterization of CRISPR/Cas9-generated *SlBASL* mutants**

(A) Confocal images of mature true leaves from M82 and *slbasl-cr#4* mutant. Stomata in purple, arrowheads indicate stomatal clusters. Scale bars: 30 $\mu$m.

(B) Confocal images of abaxial cotyledon epidermis from two additional, independently derived *slbasl* mutant lines. Arrows point to stomatal clusters.

(C) Two examples of stomatal pairs arising from fate errors in slbasl-cr#4; DIC images of same cotyledon at 1 and 3 dpe; stomatal precursors green, and stomata purple.

(D) Single found example of stomatal pair arising from fate errors in slbasl-cr#4; DIC images of same cotyledon at 1 and 3 dpe; stomatal precursors green, and stomata purple.

(E) Full characterization of CRISPR/Cas9-induced indels at the 4 gRNA sites in *slbasl-cr#4*. PAM highlighted in blue, gRNA target sequences in red. Deletions are marked by dashes; arrowhead marks a large insertion at gRNA4.

(F) Predicted protein sequences of SlBASL and *slbasl-cr#4*. Major domains of conservation denoted in colors. In the *slbasl-cr#4* allele, a small deletion at gRNA2 is predicted to lead to an early stop codon after 63 amino acids.

| Table S1: Number of BASL and BSLL proteins displayed in Fig. 1C | | | |
|---|---|---|---|
| Species | Clade | BASLs | BSLLs |
| *Physcomitrella patens* | Moss | 0 | 2 |
| *Selaginella moellendorffii* | Lycophyte | 0 | 2 |
| *Amborella trichopoda* | Basal angiosperm | 0 | 1 |
| *Ananas comosus* | Monocot | 0 | 1 |
| *Brachypodium distachyon* | Monocot | 0 | 2 |
| *Oryza sativa Japonica Group* | Monocot | 0 | 3 |
| *Triticum aestivum* | Monocot | 0 | 1 |
| *Zea mays* | Monocot | 0 | 6 |
| *Zostera marina* | Monocot | 0 | 2 |
| *Aquilegia coerulea* | Eudicot | 1 | 2 |
| *Arabidopsis thaliana* | Eudicot | 1 | 2 |
| *Brassica rapa* | Eudicot | 2 | 5 |
| *Capsella rubella* | Eudicot | 1 | 2 |
| *Capsicum annuum* | Eudicot | 1 | 4 |
| *Citrus clementina* | Eudicot | 1 | 1 |
| *Cucumis melo* | Eudicot | 1 | 3 |
| *Daucus carota subsp. sativus* | Eudicot | 0* | 1 |
| *Erythranthe guttata* | Eudicot | 0* | 1 |
| *Glycine max* | Eudicot | 1 | 5 |
| *Gossypium raimondii* | Eudicot | 2 | 4 |
| *Helianthus annuus* | Eudicot | 0* | 5 |
| *Lactuca sativa* | Eudicot | 1 | 2 |
| *Malus domestica* | Eudicot | 2 | 4 |
| *Manihot esculenta* | Eudicot | 2 | 4 |
| *Medicago truncatula* | Eudicot | 1 | 2 |
| *Nelumbo nucifera* | Eudicot | 1 | 2 |
| *Papaver somniferum* | Eudicot | 2 | 3 |
| *Populus trichocarpa* | Eudicot | 1 | 3 |
| *Prunus persica* | Eudicot | 2 | 2 |
| *Solanum lycopersicum* | Eudicot | 1 | 3 |
| *Theobroma cacao* | Eudicot | 1 | 2 |
| *Trifolium pratense* | Eudicot | 2 | 1 |
| *Vitis vinifera* | Eudicot | 2 | 3 |
| * BASL orthologues missing in the UniProtKB database, but annotated elsewhere; see for example Table S2 | | | |
| Genomes with two BASL homologues are domesticated species where there is evidence of recent genome duplication | | | |

| Table S2: Sequences used in Fig. S1C | | |
|---|---|---|
| Species | Gene name | ID |
| *Arabidopsis thaliana* | AtBASL | AT5G60880 |
| *Medicago truncatula* | MtBASL | Medtr2g461550.1 |
| *Populus trichocarpa* | PtBASL | Potri.015G048600.1 |
| *Solanum lycopersicum* | SlBASL | Solyc03g114770.2.1 |
| *Erythranthe guttata* | EgBASL | XP_012853883.1 (NCBI) |
| *Daucus carota* | DcBASL | DCAR_023155 |
| *Aquilegia coerulea* | AqBASL | Aqcoe7G067300.1 |

| Table S3: Genes names used in this paper | | | |
|---|---|---|---|
| Species | Gene name | UniProt ID | Gene ID |
| *Arabidopsis thaliana* | AtBASL | Q5BPF3 | AT5G60880 |
| *Arabidopsis thaliana* | AtBSLL1 | Q9LMY2 | AT1G13650 |
| *Arabidopsis thaliana* | AtBSLL2 | F4ITB8 | AT2G03810 |
| *Solanum lycopersicum* | SlBASL | A0A3Q7GGD1 | Solyc03g114770.2.1 |
| *Solanum lycopersicum* | SlBSLL1 | A0A3Q7FSN7 | Solyc03g114750.2.1 |
| *Solanum lycopersicum* | SlBSLL2 | A0A3Q7H7U4 | Solyc05g011830.2.1 |
| *Solanum lycopersicum* | SlBSLL3 | A0A3Q7FV09 | Solyc04g005290.2.1 |
| *Aquilegia coeruelea* | AqBASL | N/A | Aqcoe7G067300.1 |
| *Aquilegia coeruelea* | AqBSLL1 | A0A2G5F5L9 | Aqcoe3G377700.1 |
| *Aquilegia coeruelea* | AqBSLL2 | A0A2G5EPF7 | Aqcoe3G067800.1 |
| *Brachypodium distachyon* | BdBSLL1 | I1IAZ5 | Bradi3g47127 |
| *Brachypodium distachyon* | BdBSLL2 | A0A2K2CGV6 | Bradi5g12830 |
